# Tabular foundation models outperform Random Forests in predicting environmental variables from metabarcoding data

**DOI:** 10.64898/2026.11.26.769410

**Authors:** Henning Schröder, Stefan Lüdtke, Theodor Sperlea

## Abstract

Supervised machine learning is increasingly used to predict environmental variables and biotic indices from environmental DNA (eDNA) metabarcoding data. Random Forests have become the de facto standard for this task. Tabular foundation models (TFMs) are transformers pretrained on millions of synthetic datasets that make predictions through in-context learning, without dataset-specific training or hyperparameter tuning. They now outperform tree ensembles on general tabular benchmarks. Whether this advantage carries over to metabarcoding data, which are sparse, compositional and have many more features than samples, is unclear. Here, we extend a published benchmark of environmental metabarcoding datasets with four TFMs (TabPFN v3, TabPFN v3.5, TabPFN v3.5-Fast and TabICL). We compare them with Random Forests, Gradient Boosted Trees and other conventional models across eleven regression and eight classification tasks from aquatic habitats, combining several feature-selection methods and two input representations. TFMs were the top-ranked approaches overall. In regression, the best TFM, TabPFN v3.5, outperformed the best conventional model on nine of ten datasets with increases in R² between 0.04 and 0.17. In classification, gains were small (ΔF1 ≤ 0.06), as several tasks were already saturated. TFM performance was largely insensitive to whether raw counts or relative abundances were used, and benefited at most from light rarity or variance filtering. These gains came at the cost of higher computational cost at inference time and, thus, longer runtimes. We conclude that TFMs are a strong default for predicting environmental variables from metabarcoding data, and we provide practical recommendations for their use in biomonitoring.

## 1. Introduction

Over the last twenty years, DNA metabarcoding has become a workhorse for characterising microbial and other community composition in environmental samples. Because community composition responds to environmental conditions, metabarcoding data can also serve as an indicator of ecosystem state, and their integration into routine biomonitoring has repeatedly been called for (Pawlowski et al., 2021; Cordier et al., 2021). Supervised machine learning (ML) is the main tool for this purpose and is now regularly used to infer environmental variables and biotic indices from metabarcoding data in scientific contexts. Applications range from groundwater geochemistry (Smith et al., 2015), soil quality (Hermans et al., 2020) and the physico-chemistry and surrounding land cover of lakes (Sperlea et al., 2021, 2022a) to the ecological status of marine benthos (Cordier et al., 2017, 2018; Frühe et al., 2021; reviewed in Ghannam & Techtmann, 2021) and the identification of whole-community dynamics (Sperlea et al., 2026).

Across these studies, ensembles of decision trees such as the Random Forest (Breiman, 2001) have repeatedly performed well. This reflects a good fit between model and data: Random Forests tolerate high dimensionality and sparsity and require little preprocessing while performing well for small, tabular data, which sets them apart from linear models, support vector machines and deep learning models (Grinsztajn et al., 2022). In addition, feature importance measures (Breiman, 2001; Strobl et al., 2007) make them reasonably interpretable and have been used to identify potential bioindicator taxa. In a recent benchmark across 13 environmental metabarcoding datasets covering regression and classification tasks, we systematically evaluated combinations of preprocessing, feature selection and supervised model and found that Random Forests on raw count tables were a difficult-to-beat default (Zschaubitz et al., 2025). Based on those results, we suggested Random Forests and other decision tree ensembles to be the good practise default ML model class for this kind of analysis.

Since then, the landscape of supervised learning on tabular data has changed substantially. For a long time, neural networks were generally outperformed by tree ensembles on tabular data (Shwartz-Ziv & Armon, 2022; Grinsztajn et al., 2022). A first indication that this might change came when Müller et al. (2022) showed that transformers pretrained on millions of synthetic datasets generated from structural causal models can perform approximate Bayesian inference on tabular data. Rather than fitting parameters to a particular dataset, the resulting tabular foundation models (TFMs) perform in-context learning: at prediction time, the entire training set is passed to the pretrained network as context, and predictions for test samples are obtained in a single forward pass. Their mode of operation thus differs fundamentally from that of previously available ML models.

This architectural innovation was followed by rapid model development. TabPFN (Hollmann et al., 2023, 2025) proved particularly effective on small tables, a regime that has traditionally been difficult for ML. TabPFN v2.5 extended it to datasets with up to about 50,000 samples and 2,000 features (Grinsztajn et al., 2025), and methods for interpreting its predictions with Shapley values have become available (Rundel et al., 2024). Most recently, TabPFN v3.5 achieved top rankings on public benchmarks such as TabArena (Erickson et al., 2025) and introduced a faster variant for settings with limited computational resources (Jäger et al., 2026). In parallel, other TFMs have emerged, notably TabICL (Qu et al., 2025, 2026), whose model weights and pretraining procedure are openly available under a licence that also permits commercial use, as well as, more recently, models such as Kumo-Tabular (NVIDIA, 2026) and LimiX-2 (Zhang et al., 2026).

Although TFMs outperform most other ML models on tabular data in general, it is not obvious that this advantage carries over to environmental metabarcoding. Metabarcoding datasets are typically small, compositional and very sparse, and have many more features than samples (Gloor et al., 2017; Cao et al., 2021). These properties differ markedly from both the synthetic data used to pretrain TFMs and the datasets on which they have been benchmarked so far. Consistent with this uncertainty, a recent study of Baltic Sea 16S and 18S rRNA metabarcoding data found that Random Forests performed comparably to TFMs (Bell et al., 2026).

Here, we extend the benchmark of Zschaubitz et al. (2025) with four TFMs, namely TabPFN v3, TabPFN v3.5, TabPFN v3.5-Fast and TabICL, using the same evaluation protocol. We ask (i) whether TFMs outperform tree ensembles in predicting environmental variables from metabarcoding data, (ii) how feature selection and (iii) the representation of the data as raw counts or relative abundances affect their performance, and (iv) at what computational cost. We find that TFMs outperform conventional models in regression tasks, with gains in R² of up to 0.17, and perform comparably to the best conventional models in classification tasks. Unsupervised rarity and variance filters yield the best performance for all models, whereas supervised univariate feature selection degrades it, and the choice between raw counts and relative abundances has no significant effect on TFM performance. These gains come at the cost of longer runtimes, which can, however, be substantially reduced by GPU acceleration. We therefore suggest TFMs as the default model class for supervised ML on environmental DNA metabarcoding data.

## 2. Methods

### 2.1 Datasets and tasks

We re-used the regression and classification tasks from the benchmark of Zschaubitz et al. (2025), which contains environmental metabarcoding datasets of bacteria, eukaryotic microbes and fish spanning aquatic, soil and anthropogenic habitats, with ASV-level count tables as predictors and environmental variables (e.g. temperature, salinity, pH, nutrient concentrations, habitat category) as targets. In this study we report results for ten regression and eight classification tasks; the full list of datasets used in this study, target variables, sample sizes and feature counts is given in **Table 1**. No changes were made to the target variables, train/test splits or cross-validation scheme relative to the original benchmark. All tasks were evaluated with two input representations: raw ASV counts (absolute) and relative ASV abundances (relative).

**Table 1:** Overview over the datasets present in the benchmark. See Zschaubitz et al. (2025) for details of dataset preparation.

| Dataset | Description | Target region | Target feature | # samples / # features | Reference |
|---|---|---|---|---|---|
| PRJDB5465_Arc | Global subseafloor | 16S rRNA | water depth/aerobicity | 299 / 7925 | Hoshino et al. (2020) |
| PRJDB5465_Bac | Global subseafloor | 16S rRNA | water depth/aerobicity | 299 / 35623 | Hoshino et al. (2020) |
| bedford_basin_V6V8 | Bedford Basin water | 16S (V6-V8) | pressure/year | 693 / 378 | Robicheau et al. (2022) |
| bedford_basin_V4V5 | Bedford Basin water | 16S (V4-V5) | pressure/year | 753 / 489 | Robicheau et al. (2022) |
| boso_fish | Seawater samples from Boso peninsula | 12S (mito-chondrial) | avg. water_tem, salinity/site_name | 530 / 856 | Ushio et al. (2023) |
| wastewater_treatment | Sludge from 269 WWTPs, 23 countries | 16S (V4) | avg. temp/Continent | 687 / 66742 | Wu et al. (2019) |
| tara_oceans | Ocean water globally | 16S rRNA | depth/biome type | 139 / 35651 | Sunagawa et al. (2015) |
| bog_lakes | 8 bog lakes, N. Wisconsin | 16S (V4) | temp/Depth | 790 / 6208 | Linz et al. (2017) |
| eu_lakes_allsamples | 232 European lakes | 16S (V2-V3) | pH/- | 232 / 315733 | Boenigk et al. (2018), Bock et al. (2020), Nuy et al. (2020) |
| eu_lakes_allfeatures | 86 European lakes | 16S (V2-V3) | Mg/- | 86 / 315733 | Boenigk et al. (2018), Bock et al. (2020), Nuy et al. (2020) |

### 2.2 Models and feature selection

As a panel of conventional ML models, we chose Random Forests (Breiman, 2001), gradient boosting and Lasso for regression, and Random Forests, AdaBoost, support vector machines and Gaussian Naive Bayes for classification (default hyperparameters except for max_depth=5 for Random Forest classifiers). We added four TFMs: TabPFN v3 (Prior Labs Team, 2026), TabPFN v3.5 and its Fast variant TabPFN v3.5-Fast (Jäger et al., 2026), all run via the tabpfn Python package (version 9.2.0), and TabICL v2 (Qu et al., 2026; tabicl package version 2.1.1). All TFMs were used in their default configuration (n_estimators = “auto” for TabPFN, n_estimators = 8 for TabICL), without hyperparameter optimisation or fine-tuning, and without scaling or encoding of the input, as recommended by the developers.

Feature-selection methods comprised two unsupervised filters and two supervised approaches. The rarity filter retained ASVs reaching a relative abundance of at least 0.1% in at least 1% of training samples. The variance filter (VarianceThreshold) removed features with a training-set variance below 0.8 for raw counts and below 10⁻⁶ for relative abundances. The Fast Correlation-Based Filter (FCBF; Yu & Liu, 2003) discretised features and continuous targets into ten quantile bins and selected features whose symmetrical uncertainty with the target exceeded 0.01, removing features more strongly associated with an already selected feature than with the target. Univariate selection (GenericUnivariateSelect) retained the 100 features with the highest F-statistic, mutual information or χ² score. For all filters, at most 2,000 features were retained, ranked by their respective score (prevalence, variance or symmetrical uncertainty). All feature-selection steps were fitted on the training folds only.

### 2.3 Evaluation and statistics

Model performance was evaluated using the same cross-validation scheme as in Zschaubitz et al. (2025). For regression, we report the coefficient of determination (*R*^2^) on held-out test folds; for classification, we report macro-averaged F1. For each dataset, all combinations of model, feature-selection method and input representation were ranked by mean test performance (rank 1 = best) and, separately, by mean total runtime (rank 1 = fastest), and average and standard deviations of the ranks were calculated across datasets.

To assess the effect of feature selection independently of the model, we ranked the five feature-selection methods available for all models within each combination of dataset, model and input representation, averaged these ranks per dataset, and re-ranked them per dataset. We tested for differences between methods with Friedman tests, separately for TFMs and conventional models, followed by Nemenyi post-hoc tests (α = 0.05), and visualised the results as critical-difference diagrams (Demšar, 2006).

To assess the effect of converting counts to relative abundances, we compared R² between relative and absolute input for each model family, using rarity-filtered configurations only (one configuration per dataset and model). We used paired two-sided Wilcoxon signed-rank tests across datasets (n = 10) and corrected for multiple testing with the Holm method. As a sensitivity analysis, we repeated the comparison with negative R² values set to zero. Lasso was excluded from this comparison because it yielded R² < 0 on raw counts for 8 of 10 datasets.

All statistical analyses were performed in R (version 4.6.2; R Core Team, 2026) using ggplot2 (v.4.0.3, Wickham, 2016), patchwork, (v.1.3.2, Pedersen, 2025), ggrepel (v.0.9.8, Slowikowski, 2026), reshape2 (v.1.4.5, Wickham, 2007).

### 2.4 Runtime

Runtimes were measured as wall-clock time on an AMD Ryzen 7 7735HS with 16 cores and 64GB RAM and decomposed into the pipeline steps data loading, input and target transformation, fitting and application of the feature selector, model fitting, prediction on the test set, and metric computation and saving. To assess the effect of hardware, we re-ran all TFM configurations with relative input for four datasets (bedford_basin_V6V8, boso_fish, eu_lakes_allsamples and wastewater_treatment) on both CPU (AMD EPYC 7663 56-Core Processor with 8 allowed of 224 total cores and 512 GB RAM) and GPU (NVIDIA RTX 3090 with 24 GB VRAM). We report the speed-up as the median ratio of CPU to GPU runtime per model and dataset. Conventional models were run on CPU only.

### 2.5 Code and data availability

All code used to run TabPFN within the benchmark framework, together with scripts to reproduce the figures of this manuscript, has been added to the public repository accompanying the original benchmark at https://git.iow.de/sperlea/microbiome_ML_benchmark. Per-dataset, per-fold raw performance values and the resulting summary tables are deposited at 10.6084/m9.figshare.34993612. The datasets themselves are publicly available as described in Zschaubitz et al. (2025).

## 3. Results

### 3.1 Tabular foundation models outperform conventional models at the cost of runtime

In order to compare model performances and CPU runtimes across datasets and to control for dataset- and target-variable-specific systematic effects, we ranked all combinations of model, feature-selection method and input representation per task, with lower ranks attributed to higher performance and lower runtime. The results of comparing models across 11 datasets (Table 1) show a trade-off between ranked runtime and ranked performance for both regression and classification tasks (Fig. 1, Fig. S1). In regression, TFMs consistently outperformed Random Forests and the other machine learning methods, the best current models being TabPFN v3.5 and its Fast variant, followed by TabPFN v3 and TabICL (Fig. 1A, Fig. 1). In classification, TFM configurations again occupied the top performance ranks, but the separation from conventional models was less pronounced: the performance ranks of TFM configurations overlapped with those of the best Random Forest configurations (Fig. 1B). The higher performance of TFMs came at the cost of longer runtimes in both task types; this was more pronounced in classification, where every TFM configuration ranked behind every conventional configuration in total runtime (Fig. 1B).

**Fig. 1.**
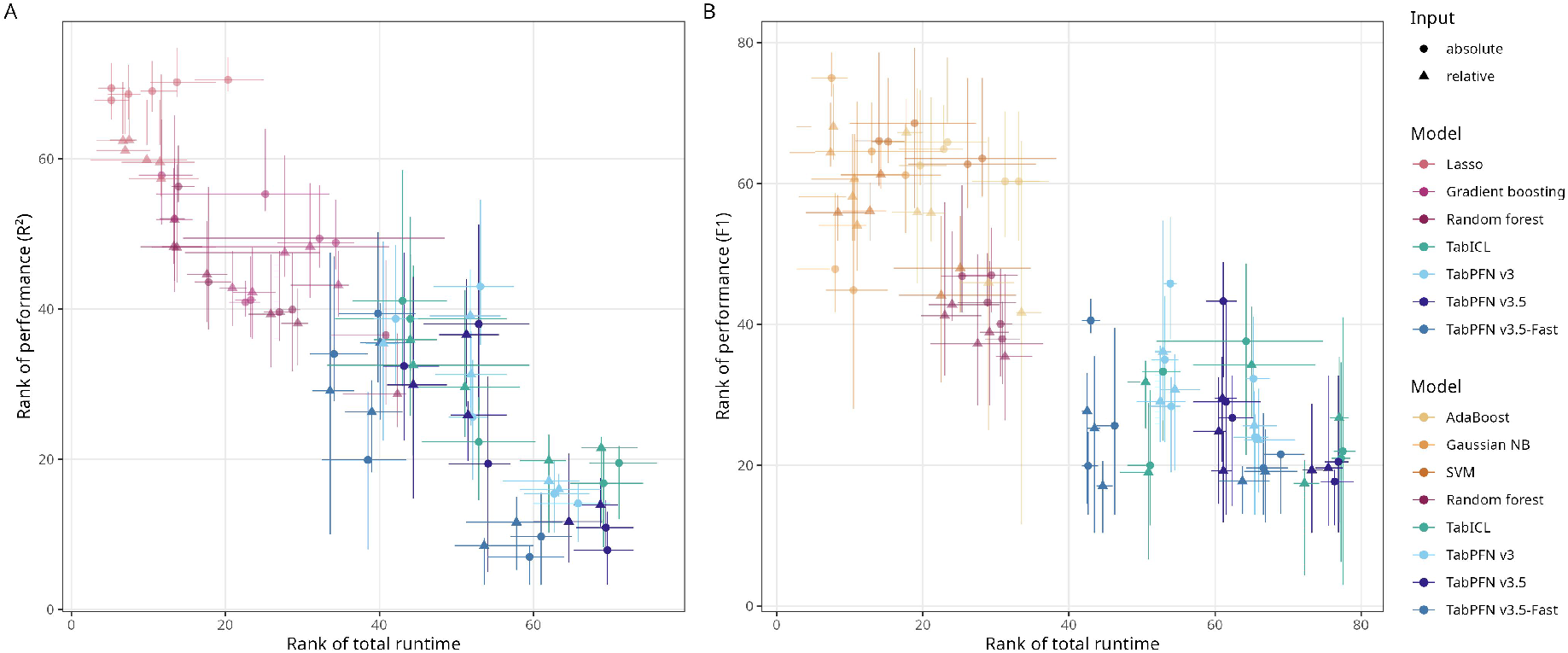
There is a trade-off between ranked performance and ranked CPU runtime in machine learning approaches in regression (a) and classification (b) tasks and TFMs generally outperform the other ML approaches. Bars represent the full range in rank.

### 3.2 Performance gains are large in regression and small in classification

In contrast to the classification tasks, TFMs bring a clear improvement over non-TFM models in regression tasks (Fig. 2). Comparing the best TFM configuration with the best non-TFM configuration for each dataset, TFMs achieved a higher R² on 9 of the 10 regression datasets for which both model groups were evaluated, with increases in R² of between 0.04 (eu_lakes_allfeatures) and 0.17 (PRJDB5465_Arc; Fig. 2A). The exception was tara_oceans, the dataset with the lowest overall predictability, on which a Random Forest reached an R² of approximately 0.32, whereas no TFM exceeded 0.16. We also observed occasional failures of individual TFM configurations; on PRJDB5465_Bac, for example, TFMs using F-statistic feature selection yielded R² values close to zero, while other configurations of the same models performed well (Fig. 2A).

**Fig. 2.**
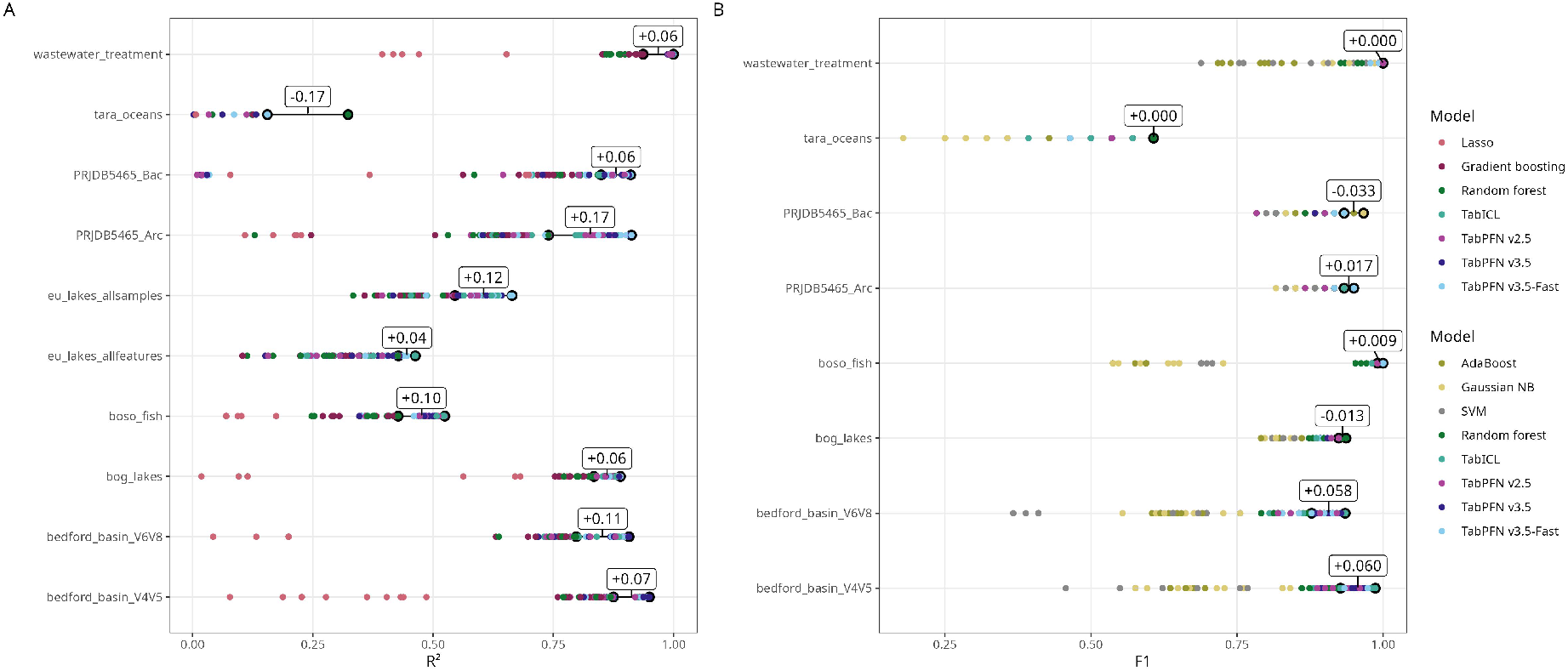
TFMs represent a clear improvement in performance in regression tasks (A) and a slight improvement in classification tasks (B). Best TFM and non-TFM model are highlighted with a black circle; numbers indicate distance between the highlighted performances in R² (A) and F1 (B). Only performance values >0 are shown.

In classification, differences between the best TFM and the best non-TFM configuration were small (ΔF1 ≤ 0.06), and on four of the eight datasets a non-TFM model performed as well as or better than the best TFM (Fig. 2B). Several classification tasks were close to the ceiling of the metric, with the best macro-F1 exceeding 0.9 on all datasets except for tara_oceans, leaving little room for improvement.

### 3.3 Rarity filtering performs best and input representation matters little, independent of model class

Depending on the dataset and the method, feature selection reduced the number of features by a factor of between 1.2 and approximately 9,100 (Fig. 3). To assess whether the choice of feature-selection method affects performance independently of the model, we ranked the five selection methods available for all models within each combination of dataset, model and input representation, and compared their mean ranks across datasets for the regression task (Fig. S2). The choice of method significantly affected performance for both TFMs (Friedman χ² = 20.3, df = 4, p < 0.001) and conventional models (χ² = 18.4, df = 4, p = 0.001; n = 10 datasets each), and the ranking of methods was the same for both model classes. The unsupervised rarity filter achieved the best mean rank for TFMs (1.70) and conventional models (1.50), followed by the variance filter (2.05 and 2.25). Supervised univariate selection based on F-statistics, together with FCBF, ranked lowest. In post-hoc comparisons (Nemenyi test, α = 0.05), the rarity filter performed significantly better than F-statistic-based selection in both model classes, whereas rarity and variance filtering did not differ significantly from each other.

**Fig. 3.**
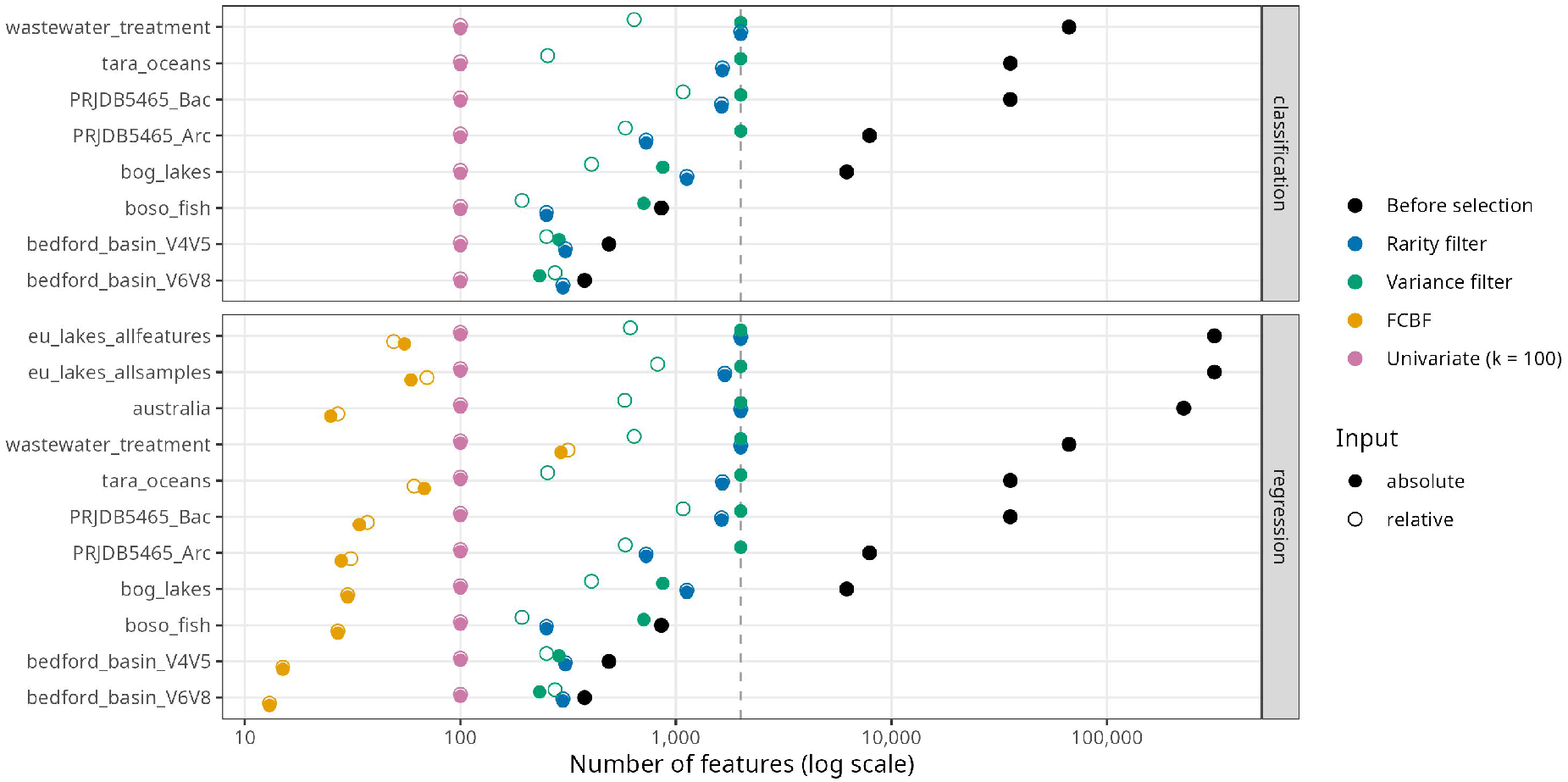
Number of features (ASVs) before and after feature selection for each dataset in the classification (top) and regression (bottom) tasks. The dashed line marks the cap of 2,000 features applied to the rarity and variance filters. Univariate selection retained a fixed number of 100 features. Note the logarithmic scale.

With rarity filtering, the choice of input representation had no significant effect on the performance of TFMs or tree ensembles (paired Wilcoxon signed-rank tests across n = 10 datasets with Holm correction, all p ≥ 0.25; Table S1). All four TFMs showed slightly negative medians, indicating at most a marginal advantage of raw counts. Lasso was excluded from this comparison because it yielded R² < 0 on raw counts for 8 of 10 datasets, rendering differences in R² uninterpretable; this is consistent with the known sensitivity of L1-regularised linear models to unscaled features.

### 3.4 GPU acceleration narrows, but does not close, the runtime gap between TFMs and conventional models

The composition of runtime differed fundamentally between conventional models and TFMs. For TFMs, most of the runtime was spent in the inference (Fig. S3), which reflects the in-context learning paradigm of these models: the training data are passed to the pretrained network only at prediction time, so the computationally expensive forward pass is attributed to prediction rather than to fitting. For conventional models, runtime was dominated either by model fitting or, depending on the feature-selection method, by fitting the feature selector. Superimposed on these model-specific patterns, we observed pronounced dataset-specific effects. On the feature-rich datasets (e.g. eu_lakes, tara_oceans and wastewater_treatment), data loading accounted for a substantial share of runtime across most models, conventional and TFM alike. On datasets with few input features (e.g. bedford_basin and boso_fish), computation dominated. Because data loading is not accelerated by GPUs, these dataset-specific differences are also relevant for interpreting the effect of hardware on runtime.

To assess how much of the runtime disadvantage of TFMs is due to hardware, we re-ran all TFM configurations for four representative datasets on both CPU and GPU (Fig. 4). GPU acceleration reduced total TFM runtimes by a factor of 1.4 to 12.3, with the largest speed-ups on the two smallest datasets (bedford_basin_V6V8 and boso_fish: 5.6–12.3×) and the smallest on eu_lakes_allsamples (1.4–2.4×), a feature-rich dataset on which data loading accounted for a substantial share of total runtime. On GPU, the total TFM runtimes on eu_lakes_allsamples and wastewater_treatment were of the same order of magnitude as those of conventional models on CPU, and TabPFN v3 on GPU was even slightly faster than the Random Forest on wastewater_treatment. On bedford_basin_V6V8 and boso_fish, TFMs remained about one order of magnitude slower than conventional models despite GPU acceleration, although absolute runtimes stayed below 10 s per configuration. Among the TFMs, TabICL benefited strongly from GPU acceleration but remained the slowest model on all four datasets. As expected, GPU acceleration had the largest effect on the prediction step, which it accelerated 18.5- to 67-fold across datasets (Fig. 4). Even so, TFM prediction on GPU remained two to three orders of magnitude slower than prediction with conventional models. The runtime of conventional models, in contrast, is dominated by other steps, such as model fitting and data loading (Fig. S3).

**Fig. 4.**
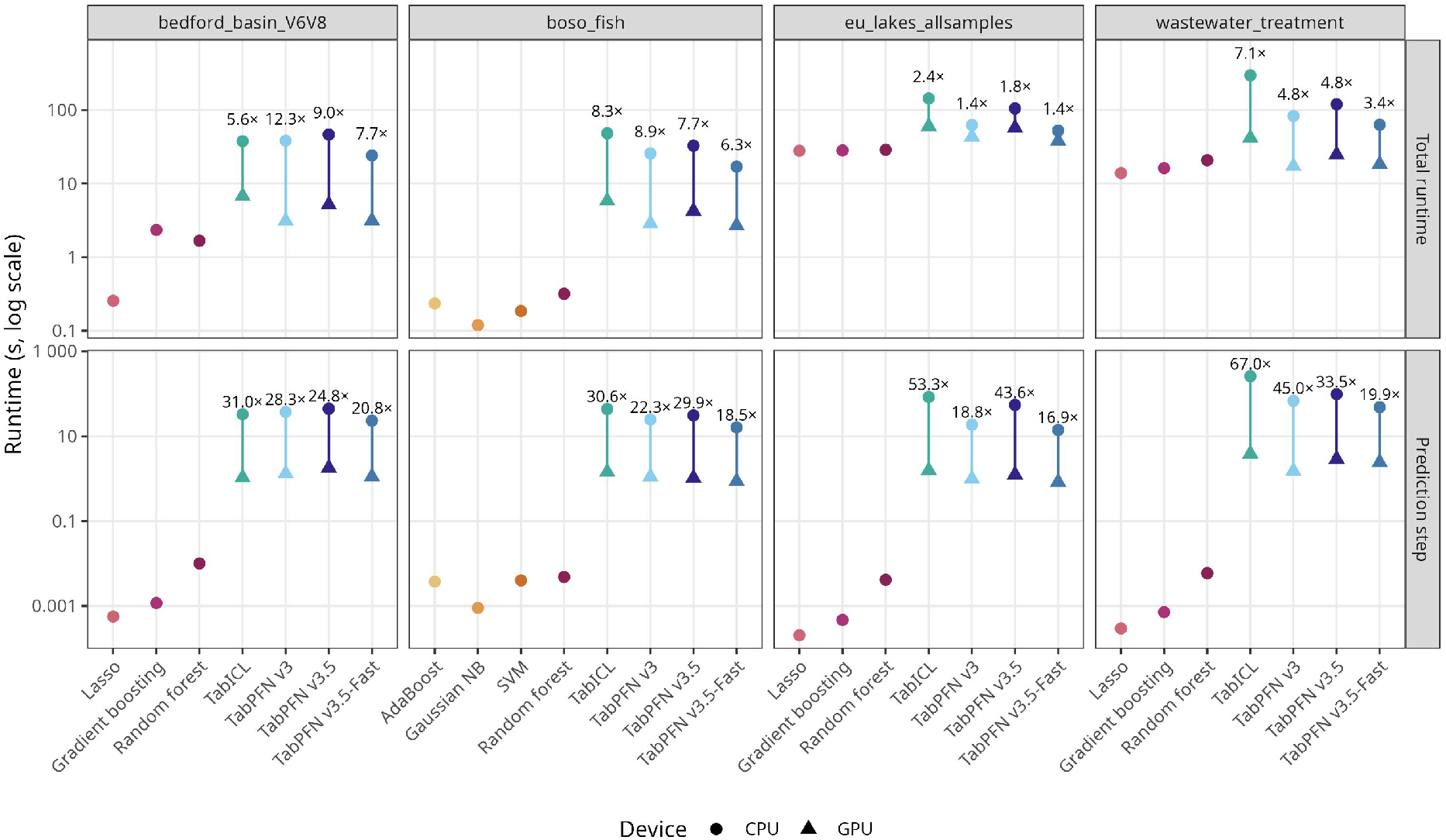
Changes in total runtime and model prediction step runtime of tabular foundation models on CPU (circles) and GPU (triangles) compared with conventional models on CPU for four datasets. Numbers give the speed-up factor (CPU/GPU runtime). Note the logarithmic scale.

## 4. Discussion

Here, we present the results of the first systematic benchmark analysis for the prediction of environmental variables from environmental DNA metabarcoding datasets that includes TFMs. Our results suggest that TFMs need to be considered as new default model class for DNA metabarcoding: They outperform conventional models in regression tasks for 9 out of 10 datasets in our benchmark and do so with an increase in R² of up to 0.17; for classification, we observe TFM performance comparable to the best conventional models, which might be due to a ceiling effect in the context of the benchmark dataset.

This result was not self-evident: metabarcoding data are sparse, compositional and have many more features than samples (Gloor et al., 2017, Cao et al., 2021), which is quite unlike the synthetic data used in the TFM pretraining process. In fact, a recent study evaluated supervised learning on Baltic Sea 16S and 18S rRNA gene metabarcoding data and reported that TabPFN perform comparably to Random Forests (Bell et al. 2026). The increase in performance we see between TabPFN v3 and v3.5 in our analysis (Fig. 1, Fig. S1) suggest that the disparity between their and our results might be due to increases in performance between version 2 and version 3.5 of TabPFN. In addition, our results highlight feature selection methods that reduce dimensionality in an unsupervised manner, thus both avoiding imposition of e.g. linear relationships between input and target and making the dataset more similar to the (presumed) pretraining data (Fig. S2). This confirms prior results regarding supervised ML (Sperlea et al., 2021, Zschaubitz et al., 2025) and is in line with the widespread practice of performing rarity filtering steps before environmental microbiome analyses (Cao et al., 2021). Furthermore, converting metabarcoding data to relative abundances had no significant effect on TFM performance in regression tasks (Table S1). This contributes to an ongoing discussion between the principled position that sequencing depth should not carry ecological meaning, since the total read count per sample is determined by the sequencing instrument rather than by the sampled community (Gloor et al., 2017; Quinn et al., 2018), and recent computational results suggesting that it might (Zschaubitz et al., 2025; Kujat et al., 2026). The results presented here suggest that, at least for TFMs, any information carried by sequencing depth is either negligible or not exploited.

There are two potential drawbacks of TFMs in environmental DNA metabarcoding when compared to conventional models. First, our results show that TFMs are generally slower than the conventional models (Fig. 1), a gap only narrowed but not overcome by running the former on GPUs (Fig. 4). However, in our estimation, in most academic settings, a slower runtime will not constitute a reason for not using a better performing model. Furthermore, it stands to be expected that further development in TFMs will decrease runtimes. Secondly, in contrast to Random Forests and other conventional supervised ML models, the training process and why it results in such a good performance must be considered opaque, potentially leading to a lower interpretability (Rudin, 2019). In practice, however, decisions by Random Forests have not been interpreted mechanistically but via feature importance scores (Breiman 2001, Strobl et al., 2007) and Shapley values (Lundberg & Lee, 2017), the latter of which are also available for TFMs (Rundel et al., 2024). This is especially important because supervised ML feature importance scores have been used to suggest metabarcoding based bioindicators (Hermans et al., 2020, Sperlea et al., 2021).

While TabPFN v3.5 outperforms other TFMs in this benchmark analysis, it is too early in the development of TFMs to advocate for a specific model. Indeed, we can see a rapid increase in performance between versions of the same model in this analysis, even if the models were published only a few years apart. After all, the Random Forest algorithm was published in 2001 (Breiman, 2001) and XGBoost, one of its most performative variants, only in 2014 (Chen et al., 2016). Indeed, at the time of writing of this manuscript, models such as Kumo-Tabular (NVIDIA, 2026), LimiX-2 (Zhang et al., 2026), Inpriora (Ant Group, 2026) and Seldon (Neuralk-AI, 2026) outperform TabPFN v3.5 on the TabArena leaderboard. We see no reason to assume that the general development of TFMs will not also result in better performance on DNA metabarcoding data. Furthermore, changes to and additions to the either the TFM pretraining or the public benchmark datasets (Erickson et al., 2025, Purucker et al., 2026) that come in the form of DNA metabarcoding datasets or at least share their characteristics will probably lead to increased performance. Thus, this field is very much in movement, but it is clear that the class of TFM produce clear gains in eDNA research over conventional models.

From an ecological point of view, the results presented here suggest that earlier estimates of how strongly communities covary with environmental variables (for example Smith et al., 2015, Hermans et al., 2020, Sperlea et al., 2021, and possibly even Bell et al., 2026) were, in many cases, lower bounds. That is to say that the community composition encodes more environmental information (in the sense of Sperlea et al., 2022b) than tree ensembles revealed and, thus, is a better indicator of ecosystem state than previously expected. Given that most datasets in the benchmark are microbial, our results support calls for integrating microbial communities as indicators of ecosystem state into monitoring programs (Pinhassi et al., 2022), and specifically for combining DNA metabarcoding with supervised ML for this purpose (Cordier et al., 2020; Frühe et al., 2021). Furthermore, TFMs like TabPFN perform a Bayesian inference that provides uncertainty estimates in the form of prediction intervals in addition to predicted values at no extra cost. In many real-world situations such uncertainty estimates are very useful, and while there are methods of preparing them from conventional models, this requires additional work and additional data and is rarely done. Finally, TFMs promise to provide a high level of performance even for small datasets, turning supervised ML a tool that is available to DNA metabarcoding datasets containing well below 100 samples.

## 5. Conclusion

Tabular foundation models are a strong new default for supervised machine learning on tabular data. In our benchmark they outperformed Random Forests and other conventional models in regression tasks and matched them in classification on DNA metabarcoding datasets, without any hyperparameter tuning. For practitioners, this translates into a few concrete recommendations. First, use a current TFM as the primary model for regression tasks, but keep a Random Forest as a baseline evaluated with the same validation scheme, since individual targets of low predictability may still favour tree ensembles. Second, reduce dimensionality with an unsupervised rarity or variance filter, as routinely applied in metabarcoding workflows, and avoid supervised univariate feature selection. Third, raw counts and relative abundances can be used interchangeably with TFMs, whereas linear models such as Lasso require scaled input. Fourth, run TFMs on a GPU where possible, which reduces runtimes by up to an order of magnitude. Fifth, report the exact model version and checkpoint, and check its license before use outside academic research. As TFMs continue to develop rapidly, we expect their advantage on metabarcoding data to grow, and their adoption to make molecular community data a more informative tool for assessing ecosystem state.

## CRediT author statement

**Henning Schröder:** Software, Formal analysis, Investigation, Writing – review & editing. **Stefan Lüdtke:** Conceptualization, Methodology, Writing – review & editing. **Theodor Sperlea:** Conceptualization, Formal analysis, Visualization, Writing – original draft, Writing – review & editing, Supervision, Project administration.

## Acknowledgements

Generative AI (Claude Opus 5.5) was used in the writing process of this manuscript. The authors take full responsibility for the content of the manuscript.

## Supplementary Figures

**Fig. S1.**
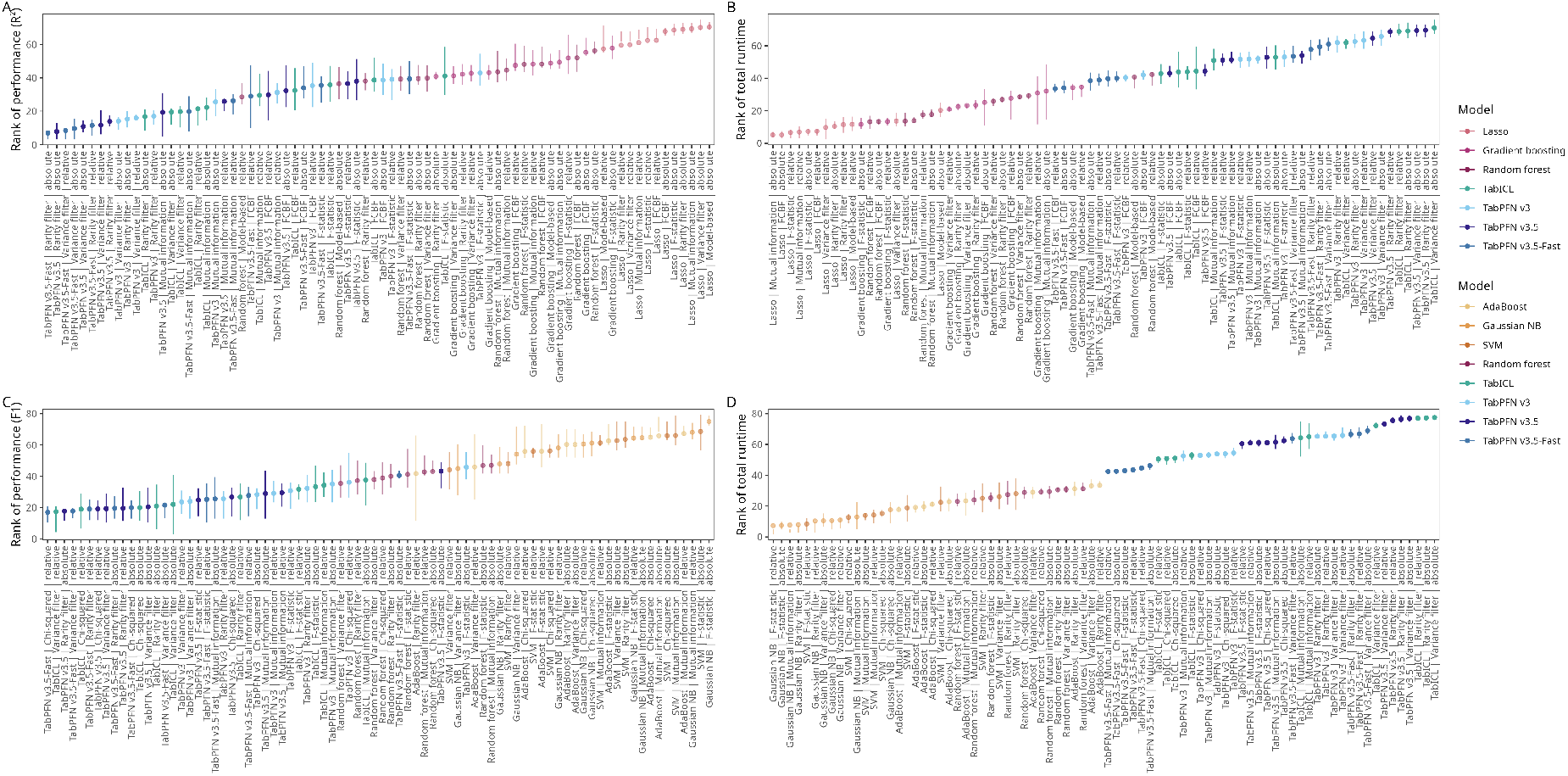
Detailed overview of the averaged ranking of performance (a, c) and total runtime (b, d) for regression (a, b) and classification (c, d) tasks. Bars represent the interquartile range of the per-dataset ranks.

**Fig. S2.**
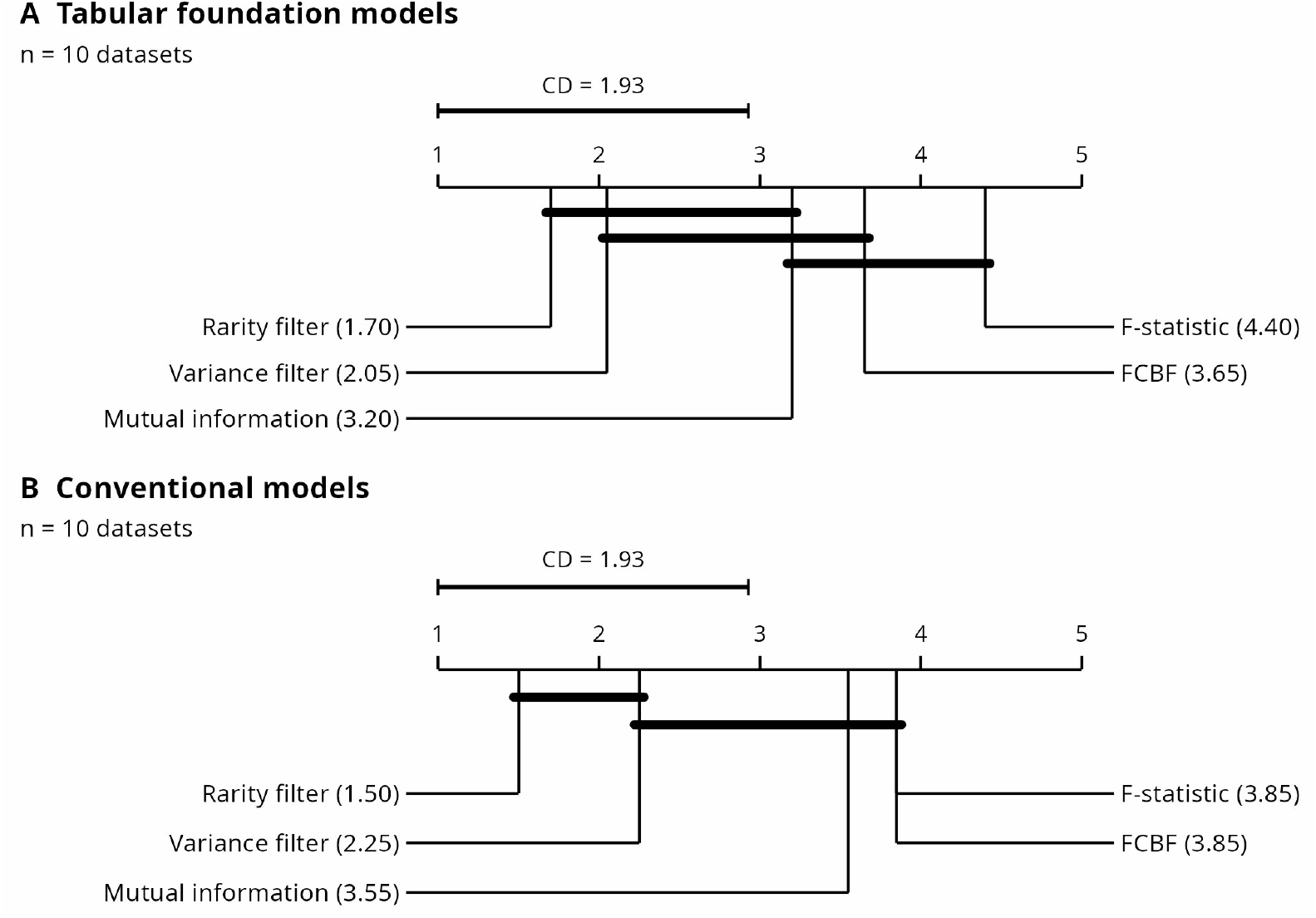
Critical-difference diagrams comparing feature-selection methods for regression tasks with tabular foundation models (A) and conventional models (B). Methods are placed by mean rank across n = 10 datasets (1 = best); numbers in brackets give mean ranks. Methods connected by a thick bar do not differ significantly (Nemenyi test, α = 0.05; CD = critical difference).

**Fig. S3.**
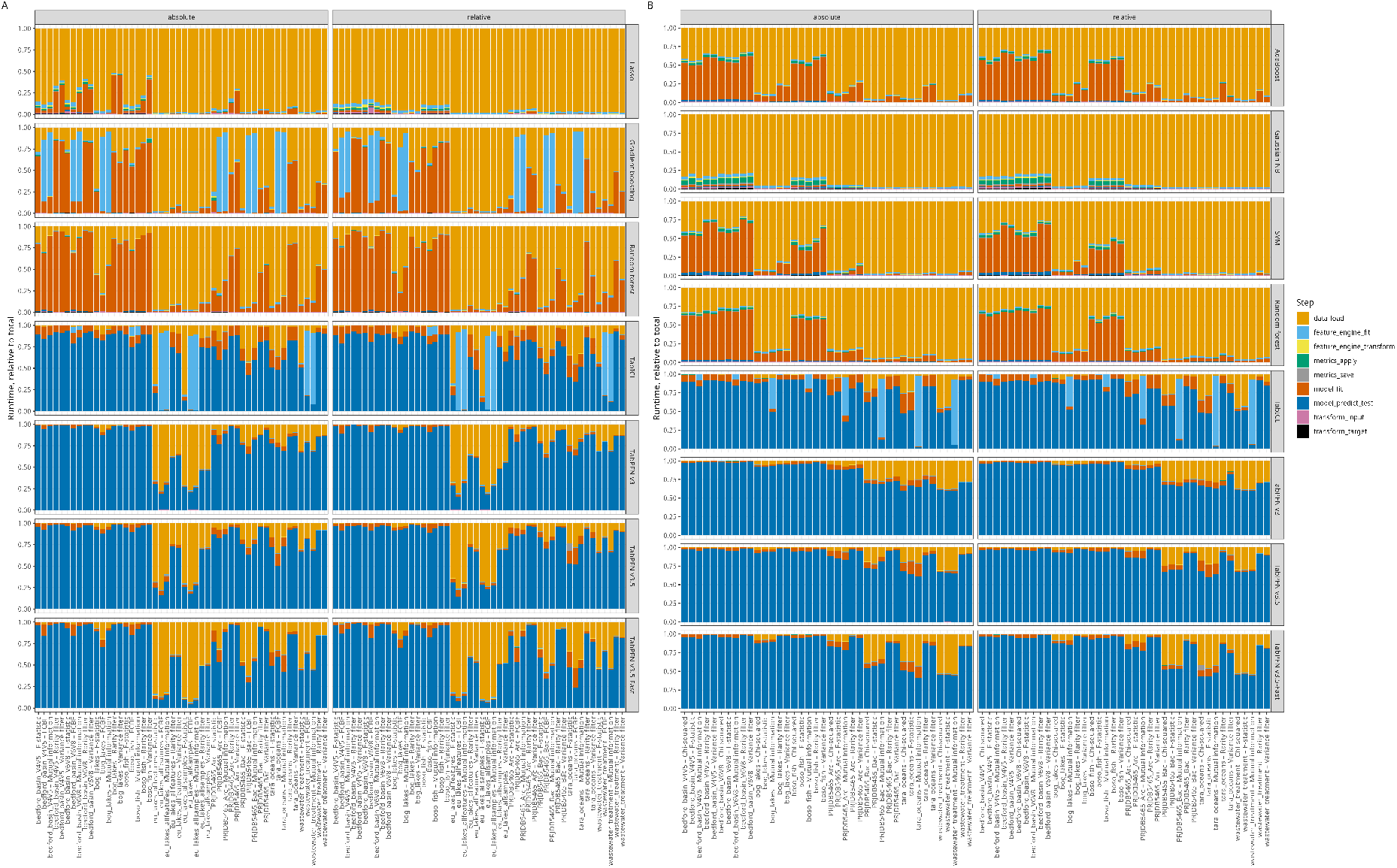
Models spend markedly different fractions of their total runtime on different steps in regression (A) and in classification tasks (B), depending on the properties of the dataset and the feature selection method used.

**Table S1.** Effect of input representation (relative abundances vs. raw counts) on regression performance with rarity filtering. ΔR² = R²(relative) − R²(absolute); median and range across n = 10 datasets; paired Wilcoxon signed-rank tests with Holm correction. Lasso was excluded (R² < 0 on raw counts for 8 of 10 datasets).

| Model | Model class | Median ΔR² | Min ΔR² | Max ΔR² | V | p | p (Holm correction) |
|---|---|---|---|---|---|---|---|
| Gradient boosting | conventional | 0.006 | -0.147 | 0.047 | 38 | 0.308 | 0.787 |
| Random forest | conventional | -0.004 | -0.045 | 0.129 | 26 | 0.919 | 0.919 |
| TabICL | tfm | -0.01 | -0.14 | 0.003 | 14 | 0.185 | 0.741 |
| TabPFN v2.5 | tfm | -0.001 | -0.059 | 0.005 | 16 | 0.262 | 0.787 |
| TabPFN v3.5 | tfm | -0.017 | -0.051 | 0.011 | 7 | 0.041 | 0.249 |
| TabPFN v3.5-Fast | tfm | -0.013 | -0.045 | 0.006 | 7 | 0.041 | 0.249 |

